# Short loop motif profiling of protein interaction networks in acute myeloid leukaemia

**DOI:** 10.1101/306886

**Authors:** Sun Sook Chung, Anna Laddach, N. Shaun B. Thomas, Franca Fraternali

## Abstract

Recent advances in biotechnologies for genomics and proteomics have expanded our understanding of biological components which play crucial roles in complex mechanisms related to cancer. However, it is still challenging to extract from the available knowledge reliable targets to use in a translational setting. The reasons for this are manifold, but essentially distilling real biological signal from heterogeneous “big data” collections is the major hurdle. Here, we aim to establish an in-silico pipeline to explore mutations and their effects on protein-protein interactions, with a focus on acute myeloid leukaemia (AML), one of the most common blood cancers with the highest mortality rate. Our method, based on cyclic interactions of a small number of proteins topologically linked in the network (short loop network motifs), highlights specific protein-protein interactions (PPIs) and their functions in AML when compared with other leukaemias. We also developed a new property named ‘short loop commonality’ to measure indirect PPIs occurring *via* common short loop interactions. This new method detects “modules” of PPI networks (PPINs) enriched with common biological functions which have proteins that contain mutation hotspots. We further perform 3D structural modelling to extract atomistic details, which shows that such hotspots map to PPI interfaces as well as active sites. Thus, our study proposes a framework for the macroscopic and microscopic investigation of PPINs, their relation to cancers, and highlights important functional modules in the network to be exploited in targeted drug screening.

## Introduction

Acute myeloid leukaemia (AML) is a complex and heterogeneous blood cancer characterised by genetic and epigenetic abnormalities together with mRNA expression changes such as amplification or deletion [1–5]. In 2016, the World Health Organization (WHO) disease classification of AML was improved by including gene expression and DNA mutation data [6]. According to this classification different sub-groups of AML are classified by specific chromosomal translocations such as t(15;17)(q22;q12), (*PML-RARA* (10%)), t(8;21)(q22;q22) *(RUNX1-RUNX1T1* (5%)), inv(16)(p13.1q22) (*CBFB-MYH11* (5%)) and 11q23 abnormalities (MLL-related (5%)), but the most commonly detected forms of AML are those with normal karyotypes (AML-NK), which account for 405–50% of patients [7–9]. Recent large-scale DNA sequencing studies have identified genetic abnormalities in AML involving several genes which have recurrent mutations in many patients [10, 11]. The most common mutations affect the amino acid sequences of FLT3, NPM1, KIT, CEBPA, TET2, DNMT3A and IDH2 [5, 12]. However, more than 31,600 mutations that affect the sequences of 7,000 proteins have been reported in AML (based on the COSMIC database [13]), most of which occur in <10% of patients and their combinatorial mutation patterns are highly variable between individual patients. Thus, although the consequences of specific mutations of certain proteins have been studied, the importance of most of the mutations in patients with AML-NK are still not identified [1, 14].

The use of reliably assembled Protein-Protein Interaction Networks (PPINs) has become common practice in the last two decades in the quest to identify biological pathways and cellular mechanisms related to newly discovered genes or disease related proteins [15–17]. In recent years, the quality and quantity of interactions shown to occur experimentally has increased substantially, particularly due to five large-scale studies using yeast-two-hybrid [18] and a panel of different protein purification/mass spectrometry methods [19-22]. Additionally, an increasing number of public protein interaction data sources [23–25] are improving proteome coverage and quality. A collaborative effort through the International Molecular Exchange (IMEx) consortium [26] is now in place to develop data formats and define curation rules to improve data integrity. However, accurate and comprehensive compilation of such heterogeneous databases is a challenging task and the currently available information is still sparse. Therefore, we are still far from having a complete proteome map for any human cell type.

Concurrently with progress in the field of PPINs, whole genome and exome sequencing projects have identified disease-related mutations and population-related variants in protein coding regions. The former, disease-related mutation information, includes data from cell lines and samples from patients, and is collated in the OMIM [27], COSMIC [13], TCGA [28] and ClinVar [29] databases. The latter, population-related variation information, is collected in dbSNP [30], 1000 Genomes [31], ExAC and gnomAD [32]. These shared resources have enabled the discovery of disease associations of mutations in the human genome [33, 34]. In establishing the impact of these variants on protein stability and function in the cell, one possibility is to evaluate the effects of disease-causing mutations on the protein 3D structure, when available. The threedimensional structure of a protein is more conserved during evolution than its linear sequence, therefore these evaluations have been used as a better proxy to predict the impacts of mutations on the biological function(s) of the mutated protein [35–37]. Unfortunately, structural determination is still challenging and therefore not available for all proteins and their interactors, making this approach also challenging on a large scale. The structure-sequence gap is still large [38] and even the use of homologous sequence(s) cannot compensate for such missing information. Therefore, the effects of many genetic variants and mutations on biological functions and the interplay among these in curated PPINs are still largely unknown. The prediction of these mutual effects is an important challenge, as it has been suggested that many complex traits are driven by large numbers of mutations, each of which has a potentially small effect on cellular function, which is propagated through a PPIN to affect biologically important core functions [39].

Different approaches have been developed to analyse such biological ‘Big Data’ effectively [40–42] and graph theory based approaches have improved our understanding of large-scale data networks in general, and PPINs [43–45] in particular. In this case, proteins are nodes and their interactions are edges. Various global and local network properties have been suggested to measure connectivity of the network and to identify sub-network modules. Previously, we defined a short-loop network motif, a cyclic interaction of a small number of proteins, as an intrinsic feature of PPINs topologically and biologically [46]. We have, in this context highlighted that short loop network motif profiling is advantageous in assessing the quality of the network and useful to extract biologically functional sub-networks.

In the study presented here, we explore the effects of genetic mutations on PPINs of AML. We focus on the mutations present in approximately half of AML patients with a normal karyotype (AML-NK) since the combined effects of disease-related mutations have not been identified yet [1, 14]. To clarify some of the underlying phenomena in this complex disease, we generated a unified large-scale human PPIN (UniPPIN) from multiple reliable sources. Mutations in AML were mapped to the UniPPIN and our short-loop network motif profiling method was applied to extract leukaemia, cancer and non-disease related mutation sub-networks. The ratio of short loops and the functional consensus across sub-networks was compared to infer features of each network. Additionally, biological functions enriched in the short loops of AML and other leukaemias were investigated. Furthermore, we propose a novel module-based concept to compare indirectly connected proteins that share protein interactions that we named ‘short loop commonality’. This has enabled us to identify functionally important protein modules that associate with AML mutated seed proteins. The commonality information has enabled us to construct a model for the threedimensional interaction of proteins, which may drive the selection of ‘hotspot’ mutations in AML patients. We show here a further use of the short loop profiling method and the combination of this with information on pathogenic variation is demonstrated useful in highlighting crucial modules that can be targeted in drug-screenings.

## Results

### Protein mutations in leukaemias reported in the COSMIC database

The aim of the study presented here is to compare the PPIN-related properties of mutated proteins that occur in different leukaemias and to predict the potential impact on the cellular functions affected. Mutations in genes which cause amino acid changes in the four leukaemias, AML, CML, T-ALL and CLL were retrieved from the COSMIC database, the most extensive resource of curated somatic mutation information about cancer, after filtering to remove synonymous mutations and single-case observations. Mutations occurring in patients who have each of the four leukaemias were analysed together to extend potential associated disease information and increase predictions of the cellular ontologies affected.

In the COSMIC database, sequencing data for 32,330 haematopoietic and lymphoid tissue samples of leukaemia patients are available: 26,127 for AML, 2,706 for CML, 1,514 for ALL and 1,983 for CLL. For each leukaemia, there are different numbers of mutated genes encoding proteins observed in at least two patients: 4,141 proteins in AML, 318 proteins in CML, 1,065 proteins in ALL and 1,802 proteins in CLL (Table 2 the first column). By comparing the proteins with mutations in each dataset, there are only 46 proteins (0.8%) (based on the UniProt Accession number) that have mutations in all four leukaemias of which half have roles as oncogenes or tumour suppressor genes, as defined in the Cancer Gene Census [47] (Table 1). Also, 27 out of 42 genes are involved in processes highlighted as the “hallmarks of cancer” [48, 49] (Table 1 and Supplementary Table S4). The predominant functions of these mutated proteins in common for all four leukaemias are: cell differentiation (GO:0030154; 28 out of 42 unique genes), system development (GO:0048731; 27 out of 42) and organelle organization (GO:0006996; 25 out of 42).

**Table 1.**
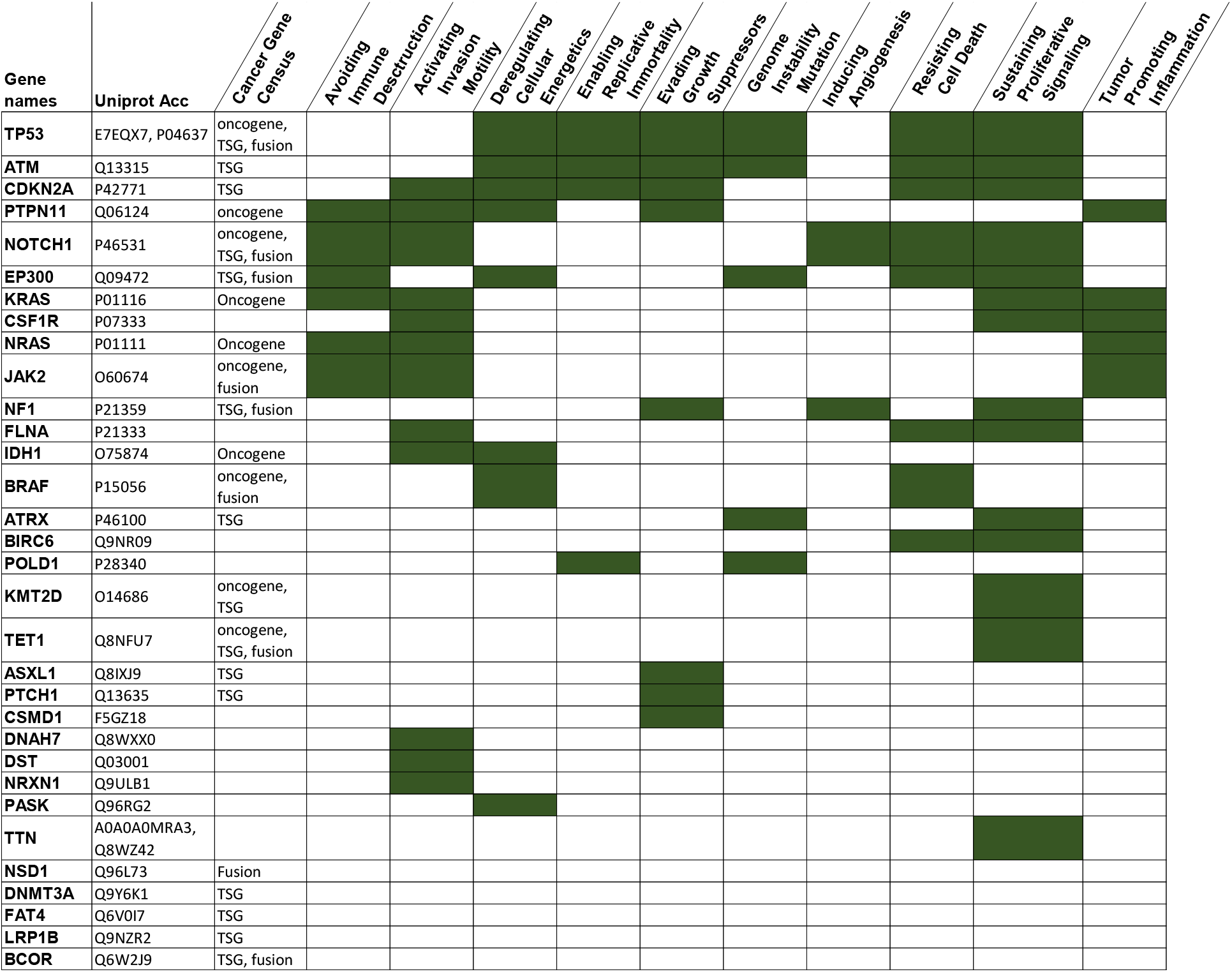
Proteins mutated in common in AML, CML, ALL and CLL and their functions in cancer. The mutated proteins in each of the four leukaemias, based on the UniProt Accession number are listed with their gene names. They were compared with the Cosmic cancer gene census information [47] (Acc: Accession number, TSG: tumour suppressor gene) and the processes related to the hallmarks of cancer [48, 49] (the gene annotations are assigned as described in [50]: shaded green). The whole list is in Supplementary Table 4.

**Table 2.**
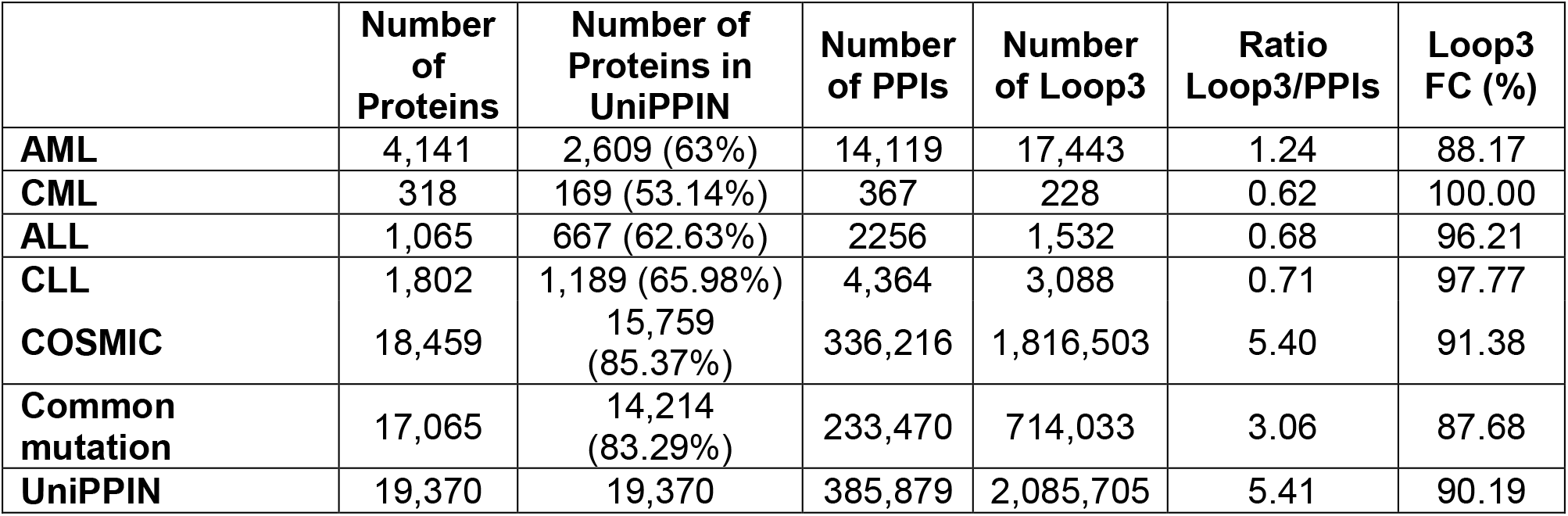
Short loop network motif (length=3) profiling for each mutation dataset. The number of short loops (length= 3) were counted and assigned with proteins having mutations in the four different leukaemias, two of which affect myeloid and two affect lymphoid cells. In each case the mutated protein was mapped onto the unified human protein interaction network (UniPPIN). Other network properties are also shown, such as the number of proteins, the number of proteins in the UniPPIN with ratios from the original number of proteins in parentheses and protein-protein interactions in the specific PPINs. In addition, the last column shows the functional consensus of the short loops, measured as the percentage of short loops having shared functions among proteins in the same loop. (FC: Functional consensus).

These terms are obtained after filtering the depth of Gene Ontology biological process terms below 3, which represent very broad terms. Although the functions of specific proteins are known, the way that mutations in these proteins affect the PPIN and the cellular processes affected is not known.

### Comparison of sub-networks by short loop network motif profiling

To investigate how proteins mutated in leukaemia can affect other proteins and pathways, their PPIN context was examined. Multiple resources of protein interaction information involving comprehensive databases and large-scale studies were employed to establish a unified human protein-protein interaction network (UniPPIN), as described in Materials and Methods. In total, there are 19,370 proteins with 385,879 interactions in the UniPPIN based on the UniProt accession number (collected on March 15^th^, 2017). Protein mutations in each leukaemia and cancer collected from the COSMIC database and non-disease related nonsynonymous single nucleotide variants (nsSNVs) from several databases were extracted as described in Materials and Methods. The somatic mutations that occur in human cancer were obtained from COSMIC [13], and the non-disease variations were obtained from a subset of dbSNP labelled as ‘common’ for variants without known pathogenic relevance, specifically of germline origin and a minor allele frequency (MAF) ≥ 0.01 in at least one major population [30]. All the proteins mutated in each leukaemia were mapped to the UniPPIN (constructed as described in Materials and Methods). Although the UniPPIN is a large-scale collection, there are gaps and more than one third of the proteins mutated in leukaemias do not map to the UniPPIN (Table 2). This is particularly true for membrane proteins, for which protein interaction data are sparse.

We showed previously that the functions of specific proteins in a large network and their local interactions with other proteins can be determined using the short loop network motif profiling method [46]. Therefore, we used this method to investigate sub-networks of proteins in the UniPPIN mutated in each of the four leukaemias and their functions. The datasets were compared by two quantitative analysis steps: 1) counting the number of short loops (length=3) that are present in each dataset and 2) measuring the consensus of the functions of proteins in each of the short loops. We also analysed short loops of length= 4 but found no significant differences with short loops of length=3 (data not shown).

The number of short loops correlates with the number of proteins and protein-protein interactions in a network (Pearson correlation score = 0.96±0.02, p-value < 4E-05) and so these were normalised by the number of protein-protein interactions, as described previously [46] (Table 2). The short loops for leukaemia-specific mutations in each of the four leukaemias were analysed. We find the normalised ratio of short loops of length 3 in AML (1.24) is significantly higher than that for all other leukaemias. It is also slightly higher (z-score= 1.32) than the normalised ratio of randomly generated PPINs (the number of random samples= 2000, the average number of proteins in random sample networks= 2602, sample mean of loop3 ratio= 0.95, sample standard deviation= 0.21, sample standard error= 0.0048) (Supplementary Figure S1). These analyses show that short loops of three proteins are particularly enriched in the AML dataset and therefore proteins mutated in AML may have more inter-connections and are possibly involved in more cooperative functions.

The overall functional enrichment of short loops was measured quantitatively by calculating the percentage of shared Gene Ontology (GO) Biological Process terms among the short loop proteins, which we define as ‘functional consensus’ [46]. This measures commonality of the functions in a loop independently of the level in the GO hierarchy and independently of the functional associations of the overall network containing the short loops. The ratio of the functional consensus in a network is calculated by the ratio of short loops having a functional consensus to all short loops 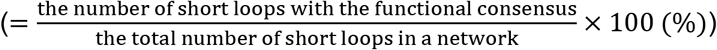. In the previous study, we highlighted that short loops in human PPINs consist of proteins with a high degree of functional consensus *(i.e.* more than 95% of short loops share at least one function) (Figure 5 in [46]). In particular 45% to 59% of the short loops in the high-confidence human PPIN [51] have a higher functional consensus ratio (>75% functional consensus). Here, we confirm these previous functional consensus analysis results [46] and find that the short loops of the human UniPPIN and all networks being analysed share at least one GO Biological Process term. Therefore, a short loop can be a ‘biological functional unit’ of the protein interaction network (Table 2). In detail, the ratios of short loops with functional consensus in the networks containing proteins mutated in the four leukaemias analysed are different and this might be due to the mutations and hence the underlying characteristics of the networks containing these mutated proteins. The short loops of length 3 in CML, ALL and CLL have a higher ratio of functional consensus than those of the PPIN containing somatic mutations in all cancers (Table 2). On the other hand, short loops in AML and non-disease related ‘common’ variation PPINs have lower functional consensus than short loops in the UniPPIN (90.19%) and COSMIC (91.38%). The lower functional consensus of short loops in AML indicates that proteins mutated in AML play roles in a wide range of biological processes. Also, GO classifications of the functions in AML short loops tend to be general and less specific (Supplementary Table S5). The biological processes in AML include metabolic processes, signal transduction and gene expression. Therefore, the results of the short loop network motif profiling of the leukaemia-specific PPINs and the functions affected by patients’ mutations reflect the complexity of the diseases, but also show that mutations in AML affect a wider range of cellular functions than those affected by mutations in the other three leukaemias or in cancer (pan-cancer analysis).

### In-depth analysis of protein mutations in AML

To determine in more detail how mutations in AML could affect cellular functions we examined how changes in the protein sequences may affect their 3D structure and functions. Based on the COSMIC database (v80, February 2017), there are more than 7,000 proteins which are affected by mutations observed in AML patients (including single patient occurrence) (Supplementary Table S6). Several large-scale studies have reported protein mutations in their patient cohorts [1, 5]. We pooled data of all AML patients to identify not only predominant mutation types in particular proteins such as FLT3 (in-frame insertion), NPM1 (frameshift insertion), CEBPA (in-frame deletion), TET2 (nonsense, in-frame deletion) and ASXL1 (frameshift, nonsense) but also the enrichment of mutations in a single amino acid position or those which localise in close vicinity in the 3D structure, defined as “mutation hotspots”. Such hotspots are composed of amino acid positions with a significantly high mutation frequency [52]. In proteins frequently mutated in AML (34 proteins having mutations observed in more than 100 patients), more than half of these proteins (19 out of 34) have hotspot mutations. Hotspot mutations account for between 50% and 99% of the mutations in these proteins. Interestingly, these mutation hotspots are located near protein binding or interaction sites (< 10 amino acid residues), when mapped on available protein 3D structures (Table 3). In the case of FLT3, Fms Related Tyrosine Kinase 3, the *FLT3-ITD* mutation is one of the most frequent primary mutations, but mutations in position D835 of the FLT3-tyrosine kinase domain (TKD) (9% of FLT3 mutations) are observed in 1328 samples (Table 3), which we analyse in more detail below. The propensity of hotspot mutations to localise to protein interacting sites could alter the functions of proteins by affecting their interactions. Therefore, we analysed protein structures further at the atomistic level.

**Table 3.** Frequently mutated proteins in AML patients and their mutation types. 7,000 proteins are mutated in AML patients according to the COSMIC (v80) database. Among them, genes with more than 100 nsSNVs are listed in the table, together with their protein names, number of mutations, their most frequent mutation types with percentage in parentheses and distance from mutation sites to protein binding or interactions sites. The information of protein binding and interaction sites are obtained from the RCSB (https://www.rcsb.org/pdb/home/home.do). Distance measures the number of amino acids difference between the sites when experimentally solved 3D structures are available. (ITD: internal tandem duplication, nsSNVs: nonsynonymous single nucleotide variants, (B): protein-ligand binding sites, (I): protein-protein Interaction sites, N/A: Not applicable (Non-hotspot mutations), PDB N/A: PDB Not available).

| Gene name | Num. of nsSNVs | Protein name | Frequent nsSNVs types | Distance (# of residue) to (B) or (I) |
| --- | --- | --- | --- | --- |
| FLT3 | 14714 | Receptor-type tyrosine-protein kinase FLT3 | ITD (84.46%), nsSNVs : D835 (9.03%) | 5 (B) |
| NPM1 | 3639 | Nucleophosmin | frameshift - insertion: W288fs* (85.54%) | N/A |
| DNMT3A | 2299 | DNA (cytosine-5)-methyltransferase 3A | nsSNVs : R882 (62.88%) | 9 (B) |
| IDH2 | 1360 | Isocitrate dehydrogenase [NADP], mitochondrial (IDH) | nsSNVs : R140 (78.01%), R172 (21.25%) | 3 (I) |
| CEBPA | 1246 | CCAAT/enhancer-binding protein alpha | insertion - In frame (30.17%), frameshift - insertion (23.6%), frameshift - deletion (23.9%) | N/A |
| TET2 | 1153 | Methylcytosine dioxygenase TET2 | nonsense (31.74%), missense SNVs (28.01%): no hotspot, frameshift - deletion (21.42%) | N/A |
| NRAS | 1093 | GTPase NRas | nsSNVs : G12 (49.77%), G13 (24.43%), Q61 (24.43%) | 0 (B) |
| IDH1 | 1077 | Isocitrate dehydrogenase [NADP] cytoplasmic (IDH) | nsSNVs : R132 (99.07%) | 3 (I) |
| WT1 | 845 | Wilms tumor protein (WT33) | frameshift - insertion (47.69%) | N/A |
| RUNX1 | 725 | Runt-related transcription factor 1 | missense SNVs (44.8%): no hotspot, frameshift - insertion (25.8%) | N/A |
| ASXL1 | 708 | Putative Polycomb group protein ASXL1 | frameshift: G646 (37.09%), nonsense (10.41%) | N/A |
| TP53 | 704 | Cellular tumor antigen p53 | nsSNVs : R248 (9.38%) | 6 (B) |
| KIT | 664 | Mast/stem cell growth factor receptor Kit | nsSNVs : D816 (57.53%), N822 (13.4%) | 6 (B) |
| MUC12 | 614 | Mucin-12 | missense SNVs (99.67%): no hotspot | N/A |
| JAK2 | 528 | Tyrosine-protein kinase JAK2 | nsSNVs : V617 (99.24%) | 10 (B) |
| PTPN11 | 364 | Tyrosine-protein phosphatase non-receptor type 11 | nsSNVs : E76(17.31%), A72(16.76%), D61(12.91%) | 1 (I) |
| MUC4 | 348 | Mucin-4 | missense SNVs (100%) : no hotspot | N/A |
| KRAS | 340 | GTPase KRas | nsSNVs : G12 (52.64%), G13 (23.24%), Q61 (9.7%) | 1 (B) |
| TTN | 313 | Titin | missense SNVs (97.12%) : no hotspot | N/A |
| SRSF2 | 254 | Serine/arginine-rich splicing factor 2 | nsSNVs : P95 (86.51%) | 0 (B) |
| BCOR | 234 | BCL-6 corepressor | nonsense nsSNVs (28.63%), frameshift - insertion (28.63%) | N/A |
| U2AF1 | 208 | Splicing factor U2AF 35 kDa subunit | nsSNVs : S34 (57.21%), Q157 (25%) | PDB N/A |
| STAG2 | 168 | Cohesin subunit SA-2 | nonsense nsSNVs (52.98%) | N/A |
| GATA2 | 163 | Endothelial transcription factor GATA-2 | nsSNVs : A318 (16.56%), R362 (14.11%), L321 (14.11%) | PDB N/A |
| EZH2 | 162 | Histone-lysine N-methyltransferase EZH2 | nsSNVs : R690 (10.74%) | 4 (I) |
| LPA | 148 | Apolipoprotein | missense SNVs (99.32%) : no hotspot | N/A |
| MUC16 | 147 | Mucin-16 | missense SNVs (98.63%) : no hotspot | N/A |
| SF3B1 | 132 | Splicing factor 3B subunit 1 | nsSNVs : K700 (37.12%) K666 (34.09%) |  |
| MUC6 | 130 | Mucin-6 | nsSNVs : P1965 (15.38%) | PDB N/A |
| GATA1 | 127 | Erythroid transcription factor | frameshift - insertion (30.71%), frameshift - deletion (27.56%) | N/A |
| PHF6 | 118 | PHD finger protein 6 | missense SNVs (42.37%) : no hotspot | N/A |
| CSF3R | 112 | Granulocyte colony-stimulating factor receptor | nonsense nsSNVs (52.67%), nsSNVs : T618(25.42%) | N/A |
| HRNR | 107 | Hornerin | missense SNVs (100%) : no hotspot | N/A |
| RAD21 | 106 | Double-strand-break repair protein rad21 homolog | nonsense nsSNVs (37.74%) | N/A |

### A protein’s short loop similarity reveals possible functional complementing roles

Short loop network motif profiling was applied to the PPIN containing proteins mutated in AML and topological and functional analyses were carried out on the short loops identified that contain proteins targeted by leukaemia mutations. Among the enriched short loops, we found some proteins do not directly interact but engage in similar short loop interactions in the sense that they engage in protein-protein interactions with the same proteins which in turn interact with each other (example Figure 1). We defined the term ‘short loop commonality’ (commonality: “The state of sharing features or attributes”:https://en.oxforddictionaries.com/definition/commonality) to describe such protein relationships. To reduce bias caused by proteins having only a few short loop interactions, we therefore defined short loop commonality proteins based on two criteria: 1) at least three short loops are shared between the proteins in a commonality relationship and 2) 95% of their short loops are in common.

**Figure 1.**
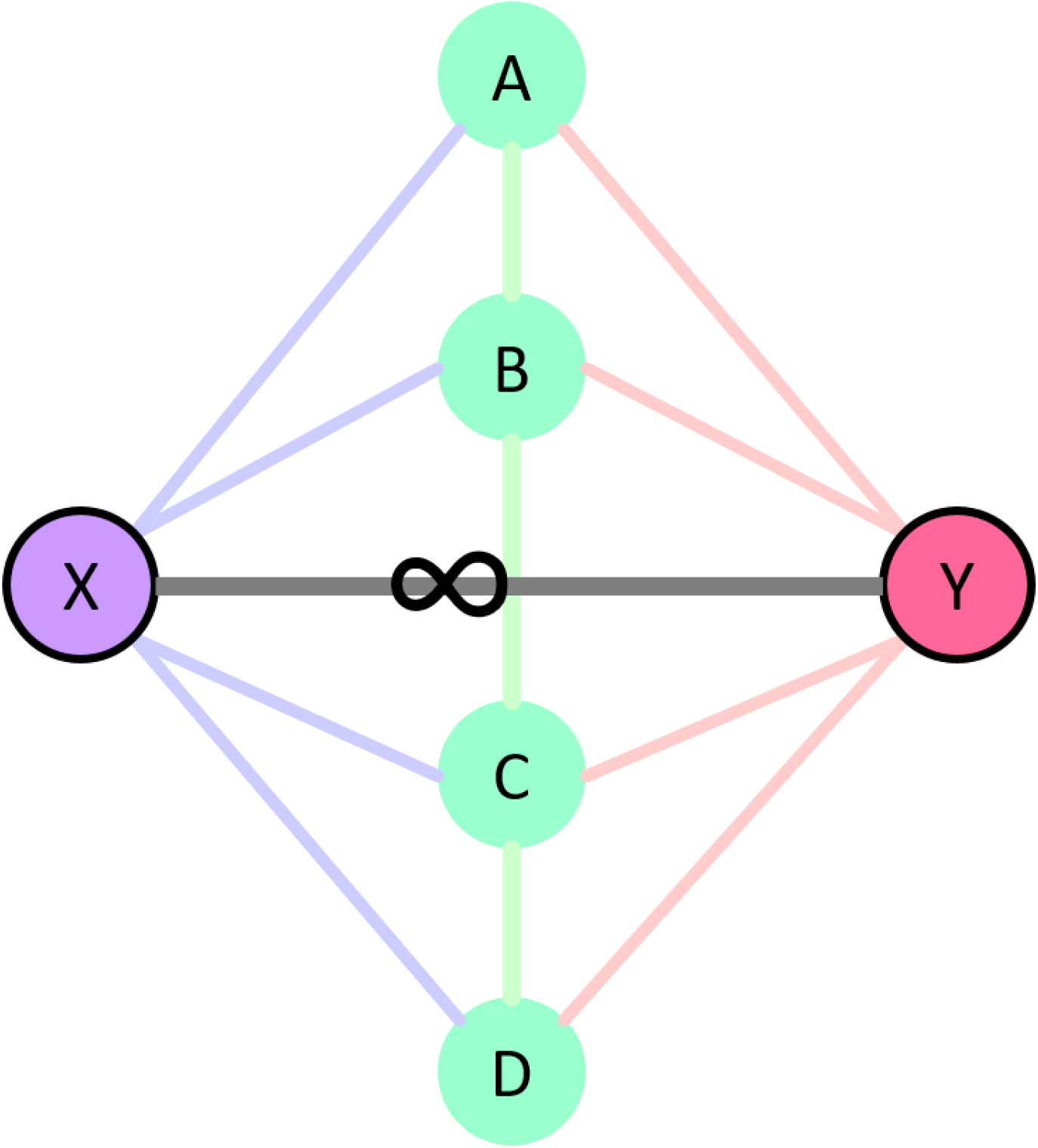
Schematic representation of short loop commonality. ‘Short loop commonality’ is defined as the association of proteins having the same or similar short loop interactions, but not involving direct interaction of the proteins themselves. For example, X forms short loops with AB, BC and CD pairs and Y also forms short loops with AB, BC and CD but there is no direct interaction between X and Y. The short loop commonality pair of X and Y is annotated with a symbol of a loop (∞).

In the network containing proteins mutated in AML, 183 proteins form 224 protein pairs which are in short loop commonality with each other (Figure 2 and Supplementary Table S7) and there are six communities or clusters of the commonality pairs involving more than 5 proteins in each. Proteins in each cluster tend to have enriched functions based on ClueGO analysis [53], such as RNA splicing, keratinization, centrosome organization and phosphatidylinositol 3-kinase (PI3K) signalling (Supplementary Table S8), which may play a role as a functional unit or “module”. Among these clusters, a cluster of 25 proteins enriched in the PI3K pathway consists of two subclusters, one with the receptor tyrosine kinase (RTK) family such as FLT3, KIT, PDGFRB, ERBB2 and MET (Figure 3 (right) and Table 3) and the other with short loop interaction partners of these RTK proteins involving PIK3R1, PTPN11, PTPRJ, CBL and CBLB (Figure 3 (right) and Table 3). These two sub-clusters are connected by a short loop commonality pair of MPL and PIK3R1 which have short loop PPIs with JAK2, SOCS1, SHC1 and PTPN11 (Supplementary Table S7). Moreover, the short loop interactions related to these RTK proteins (Figure 4 and Table 4) are enriched with Src Homology 2 (SH2) domains which are present in five out of six proteins. Thus, such PPIs with predominant functional domains lead us to the hypothesis that the functions of the receptor tyrosine kinase family are shared or overlap in the underlying short loop protein-protein interactions and therefore commonality with mutations in cancers could be enriched in such short loops.

**Figure 2.**
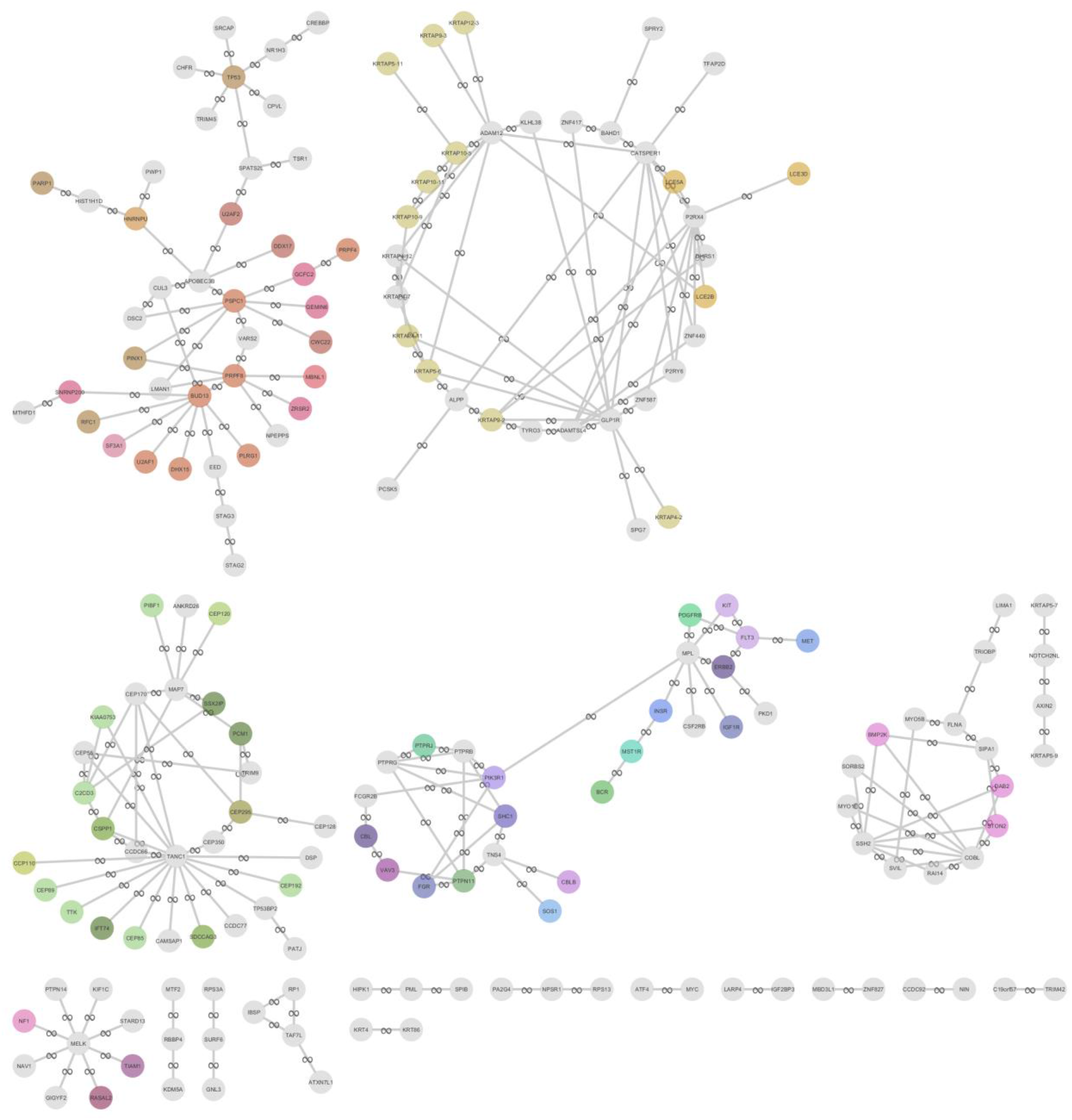
Landscape of short loop commonality among proteins with mutations in the AML PPIN. Short loop commonality, similarity of short loops between proteins, was calculated by comparing sets of protein interacting partner pairs for all proteins in the AML PPIN. In total 183 proteins (shown in light blue circles) account for 224 pairs of short loop commonality, which are annotated as line edges with a loop (∞) symbol. Functional enrichment of each cluster was measured by ClueGO [53] and nodes are coloured when proteins have enriched functional terms in the cluster. Detailed protein pairs and enriched functional terms are listed in Supplementary Table S7 and S8.

**Figure 3.**
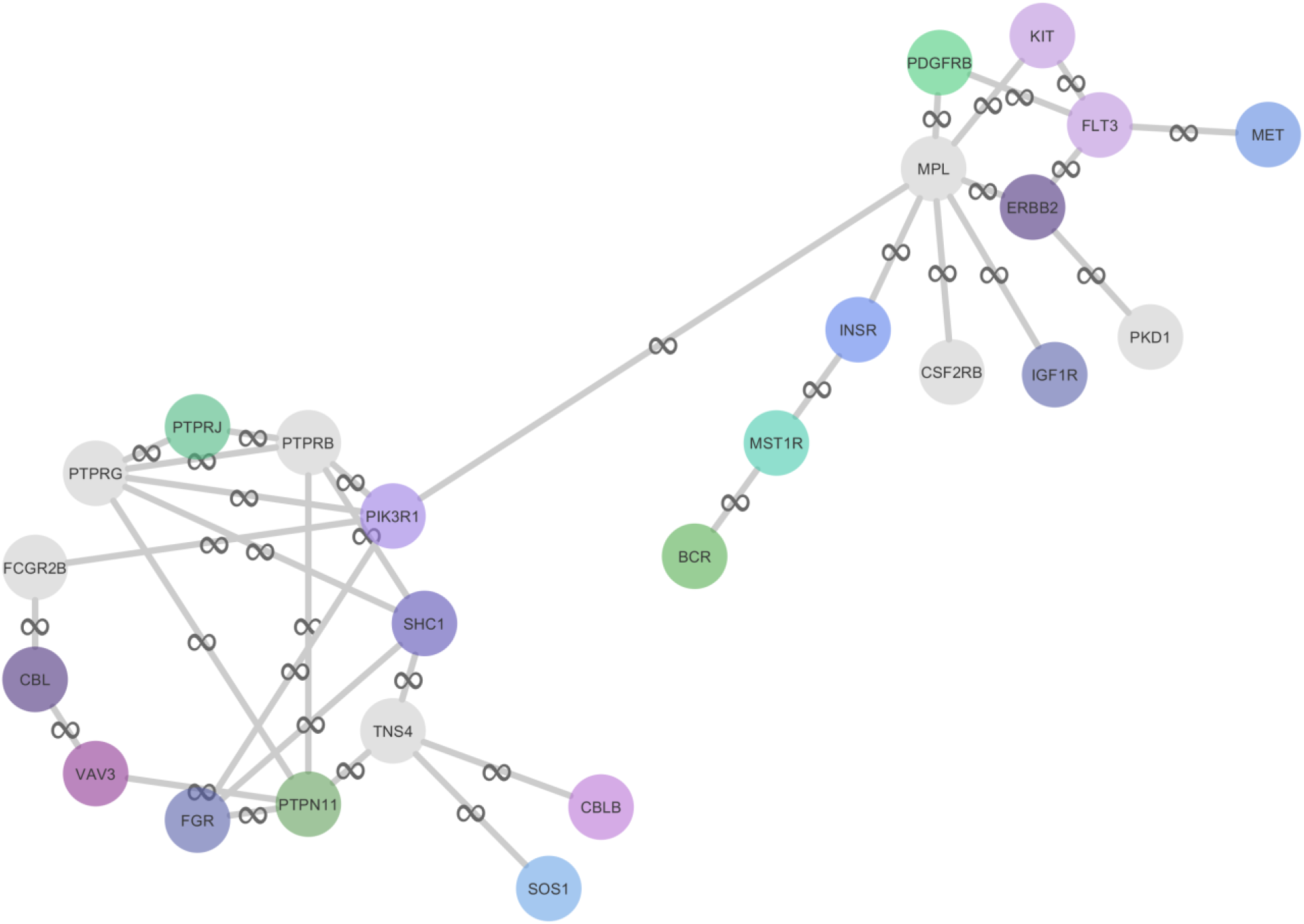
A cluster of short loop commonality in AML. The cluster with RTK proteins was extracted and proteins and their functions are annotated based on ClueGO analysis. This cluster can be divided into two subsets enriched with the RTK proteins (*e.g.* FLT3, KIT, ERBB2, PDgFrB and MET (right)) and their short loop interacting partners (*e.g.* PIK3R1, PTPRJ, PTPN11, CBL and CBLB (left)). Detailed functional terms are listed in Supplementary Table S8 and S9.

**Figure 4.**
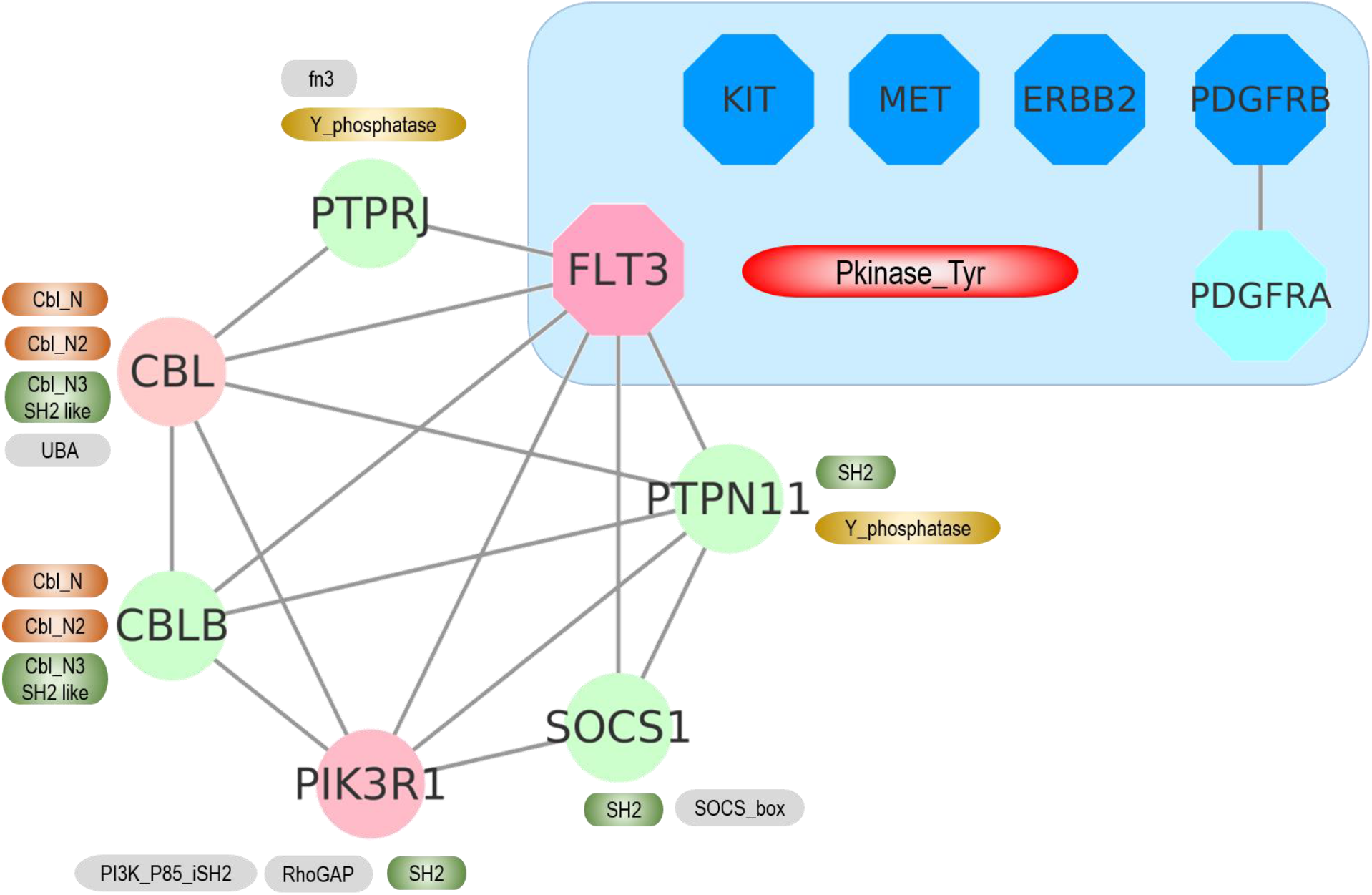
FLT3 short loop interactions and short loop commonality proteins. The left side shows that FLT3 (pink octagon) has short loop protein-protein interactions with PTPN11, PTPRJ, SOCS1, PIK3R1, CBLB and CBL proteins, annotated as circles. Their protein-protein interactions are drawn as edges. Next to each protein, their functional domains are marked in round edged boxes. The detailed domain information is in Table 4. The right upper side in the blue area shows FLT3 short loop (length=3) commonality proteins KIT, MET, ERBB2, PDGRB in blue octagons having the same short loop protein interactions as FLT3.

**Table 4.** FLT3 short loop commonality proteins, their short loop proteins and their functional domains. FLT3 has short loop commonality with 4 different protein kinases, KIT, PDGFRB, MET and ERBB2. Among them, PDGFRB is in a complex with its paralog protein, PDGRFA. FLT3 consists of 9 short loop (length=3) interactions with 6 proteins, PIK3R1, PTPN11, SOCS1, PTPRJ, CBL and CBLB. Their protein names and functional domains based on the Pfam [54] are listed.

| UniProt AC | Gene name | Protein names | Pfam domains |
| --- | --- | --- | --- |
| P36888 | FLT3 | Receptor-type tyrosine-protein kinase FLT3 | PF00047; ig [47]<br>PF07714; Pkinase_Tyr [48] |
| P10721 | KIT | Mast/stem cell growth factor receptor Kit | PF00047; ig [47]<br>PF07714; Pkinase_Tyr [48] |
| P09619 | PDGFRB | Platelet-derived growth factor receptor beta | PF07679; I-set [49]<br>PF00047; ig [47]<br>PF07714; Pkinase_Tyr [48] |
| P08581 | MET | Hepatocyte growth factor receptor | PF07714; Pkinase_Tyr [48]<br>PF01437; PSI [50]<br>PF01403; Sema [51]<br>PF01833; TIG [52] |
| P04626 | ERBB2 | Receptor tyrosine-protein kinase erbB-2 | PF00757; Furin-like [53] |
|  |  |  | PF14843; GF_recep_IV [54]<br>PF07714; Pkinase_Tyr [48]<br>PF01030; Recep_L_domain [55] |
| P16234 | PDGFRA | Platelet-derived growth factor receptor alpha | PF07679; I-set [49]<br>PF07714; Pkinase_Tyr [48] |
| P27986 | PIK3R1 | Phosphatidylinositol 3-kinase regulatory subunit alpha | PF16454; PI3K_P85_iSH2 [56]<br>PF00620; RhoGAP[57]<br>PF00017; SH2 [58, 59] |
| Q06124 | PTPN11 | Tyrosine-protein phosphatase non-receptor type 11 | PF00017; SH2 [58, 59]<br>PF00102; Y_phosphatase [60] |
| O15524 | SOCS1 | Suppressor of cytokine signaling 1 | PF00017; SH2 [58, 59]<br>PF07525; SOCS_box [61] |
| Q12913 | PTPRJ | Receptor-type tyrosine-protein phosphatase eta | PF00041; fn3 [62]<br>PF00102; Y_phosphatase [60] |
| P22681 | CBL | E3 ubiquitin-protein ligase CBL | PF02262; Cbl_N [63]<br>PF02761; Cbl_N2 [63]<br>PF02762; Cbl_N3 [63]<br>PF00627; UBA [64] |
| Q13191 | CBLB | E3 ubiquitin-protein ligase CBL-B | PF02262; Cbl_N [63]<br>PF02761; Cbl_N2 [63]<br>PF02762; Cbl_N3 [63] |

### FLT3 short loop commonality contains mutation hotspots in cancers

Mutations in kinase proteins have been extensively studied to understand cancer mechanisms [55, 56] and mutation hotspots in these proteins are observed in various cancers [57, 58]. We hypothesise that when mutated members of the RTK proteins are in a short loop commonality relationship, mutations will be in mutation hotspots in multiple cancers. By investigating these RTK proteins using mutation data from the COSMIC database, we found that FLT3 and its short loop commonality RTK proteins have frequently mutated positions or mutation hotspots in their kinase domain in various cancers (Supplementary Table S6). By analysing the ratio of the occurrence of the mutation hotspot to the total number of mutations in the corresponding RTK proteins in all cancer types (defined as Mutation Hotspot Ratio Density (MHRD), 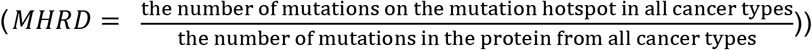, we observe this number to vary between 3–30%. All the ratios are statistically significant (p-value < 0.05, z-score > 3), assuming uniform distributions of the mutations in the protein sequences (Supplementary Table S6). Since the effect of the mutational hotspot will be dependent on its spatial position, we analysed the location specificity of mutation hotspots in the kinase domains of FLT3, and its short loop commonality proteins, in 3D structural space as well as in the linear amino acid sequences (Figure 5 and Figure 6). Figure 5 shows that the mutation hotspots are closely aligned and located near the amino acid residues, Asp-Phe-Gly (DFG) motif typical of protein kinases, known as a “gatekeeper” of protein kinase activities [59]. They are in the activation loop of the kinase domain which is located near ligand or small molecule binding sites [60]. Because of this 3D spatial closeness between mutation hotspots and functional sites, such mutations could interfere with protein-ligand interactions.

**Figure 5.**
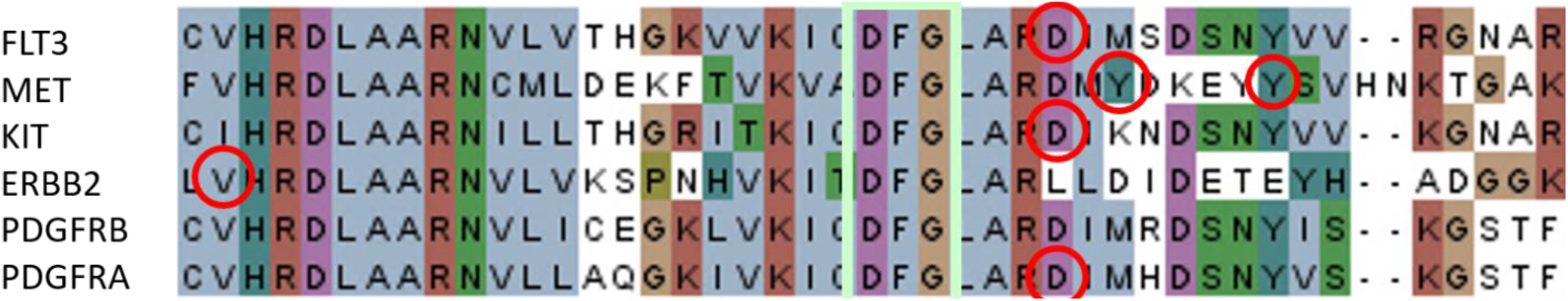
Amino acid sequence alignment of FLT3 and its commonality proteins. The protein kinase domains of each protein were aligned by T-coffee (http://tcoffee.crg.cat/) and visualized by Jalview (http://www.jalview.org/). DFG motifs are boxed in light green and the hotspot mutations of each kinase are circled in red.

**Figure 6.**
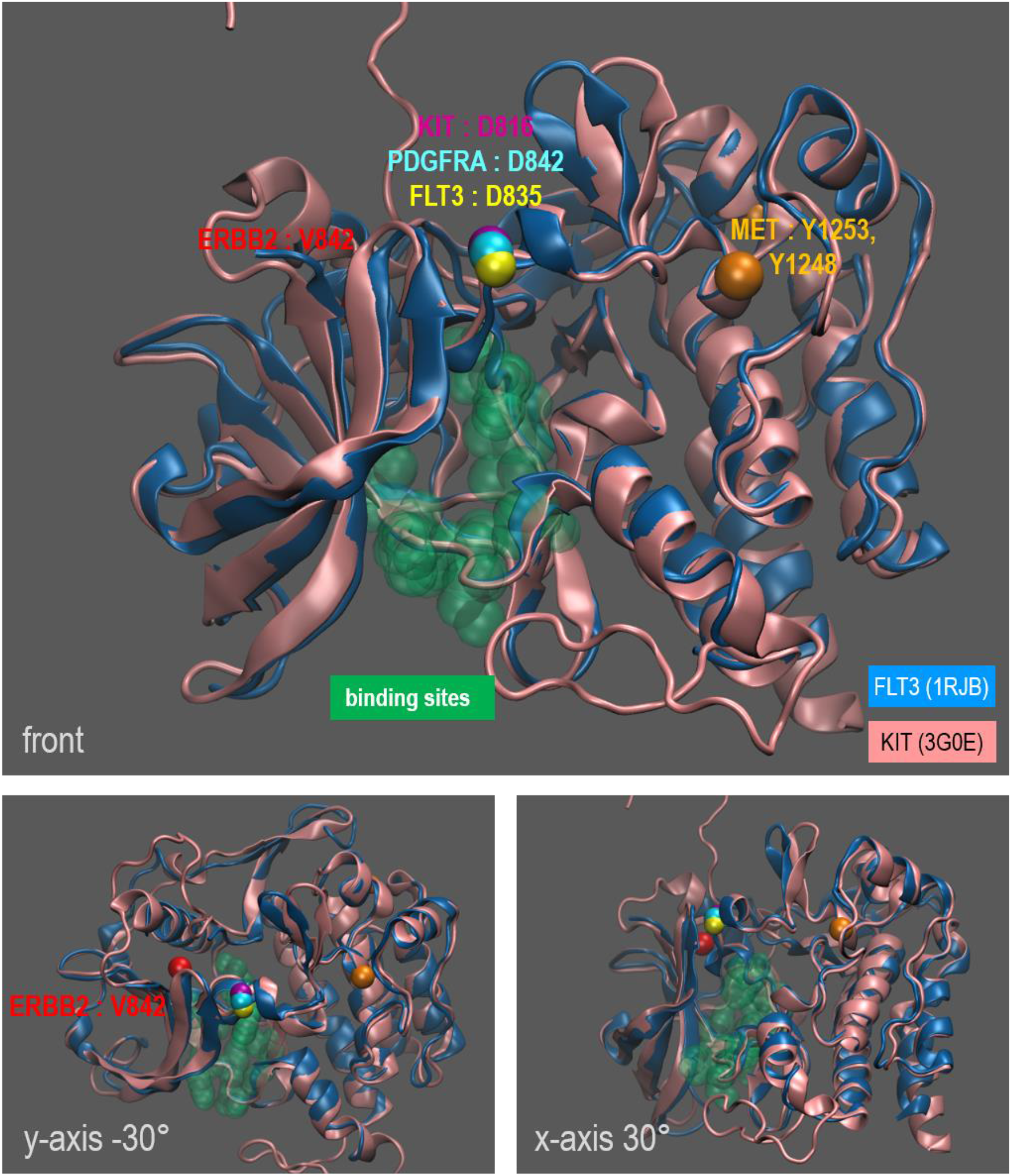
A zoomed-in view of the structural alignment of FLT3 and KIT proteins with mutation hotspots of FLT3 short loop commonality proteins in 3D space. FLT3 (1RJB) and KIT(3G0E) kinase domain structure 3D superimposition. The mutation hotspots of other FLT3 short loop commonality proteins are shown in beads representation and mapped onto this superimposition (FLT3: yellow, KIT: magenta, PDGFRA: light blue, MET: orange, ERBB2: red). To show the vicinity of mutation hotspots and the known small molecule binding site, this site is annotated on the original KIT structure and visualised as a green surface (top). Views from different angles are shown, rotating −45° in the y-axis (bottom left) and 30° in the x-axis (bottom right).

### Protein interactions of FLT3 and its short loop commonality have different interfaces for interactions with other proteins

The enrichment of cancer-related mutation sites in FLT3 and its short loop commonality proteins leads to the hypothesis that their short loop interactions with “modules” of associated proteins constrain the mutation sites that have functional effects to hotspots that affect interactions of proteins in the short loops. Among six of the interacting proteins in the short loop of FLT3 and other short loop commonality kinases such as PIK3R1, PTPN11, PTPRJ, SOCS1, CBL and CBLB we note that all except for PTPRJ have SH2 domains (Pfam ID: PF00017) or domain Cbl_N3 defined as SH2-like domains (Pfam ID: PF02762) with roles in binding to a partner protein that is phosphorylated on tyrosine [61, 62] (Figure 4). However, tyrosine residues in the RTK proteins involved are generally not found as mutated hotspots in our analyses (except for MET). Therefore, the mutation hotspots in these RTKs might affect protein-protein interaction sites rather than targeting the active site directly. To better understand the consequences that such mutations may have, we investigated the possibility that these hotspots affect residues at the interface with partner domains in the specific case of the FLT3 tyrosine kinase domain (Pfam ID: PF07714) and its partner protein domain, SH2-like domain in CBL (Pfam ID: PF02762). There is no available experimentally solved structure for the protein complex, therefore a 3D structural model was predicted for interactions of the active state of FLT3, which binds to ATP with an open conformation of the activation loop [63, 64] (known as “DFG-motif Asp-IN” and “αC Helix-IN” conformation). Also, the FLT3 mutation hotspot D835 is predicted to change the kinase domain to an active state, as has been shown to occur for the L861 mutation of EGFR [65].

The predicted models were determined by PRISM [66] and Rosetta docking [67] (see Materials and Methods; Figure 8) and show that the “active” FLT3 kinase may interact with SH2 domains *via* the middle of the kinase domain between N-and C-lobes. Interestingly, the mutation hotspot D835 is located at the interfaces of the predicted PPI models (Figure 7). This might indicate that mutation hotspots can cause a dual impact on functions of protein-small molecule binding and protein-protein interactions depending on the activation state. Furthermore, the effects of frequent mutations (D835Y, D835V, D835H; Supplementary Table S6) were predicted by mCSM (http://biosig.unimelb.edu.au/mcsm/) [68] which predicts the effect of mutations in 3D structural proteins. The predicted protein-protein affinity change (ΔΔG) upon D→ Histidine (H) or Valine (V) is slightly positive, therefore stabilising (ΔΔG > 0; average ΔΔG 0.61 and 0.12 respectively) but the change upon D→Y is slightly negative, therefore destabilising (average ΔΔG= −0.38) for the ten refined PPI models. Based on these predictions, one could argue that destabilisation of the interface is caused by this mutation. Interestingly, a previous study showed that the hotspot mutation FLT3 D835Y induces a phosphorylation site, p-Y842 [69] which is located near the predicted PPI interface (Figure 7). Thus, in summary we hypothesise that for FLT3 in an active state the hotspot mutation D835Y: 1) destabilises protein interactions of the FLT3 kinase and SH2 domain interactions; 2) exposes additional phosphorylation sites (gain-of-function role); and 3) competes for phosphorylation actively because the residue is mutated to a tyrosine and the residues of 835 and 842 are closely located at the PPI interface.

**Figure 7.**
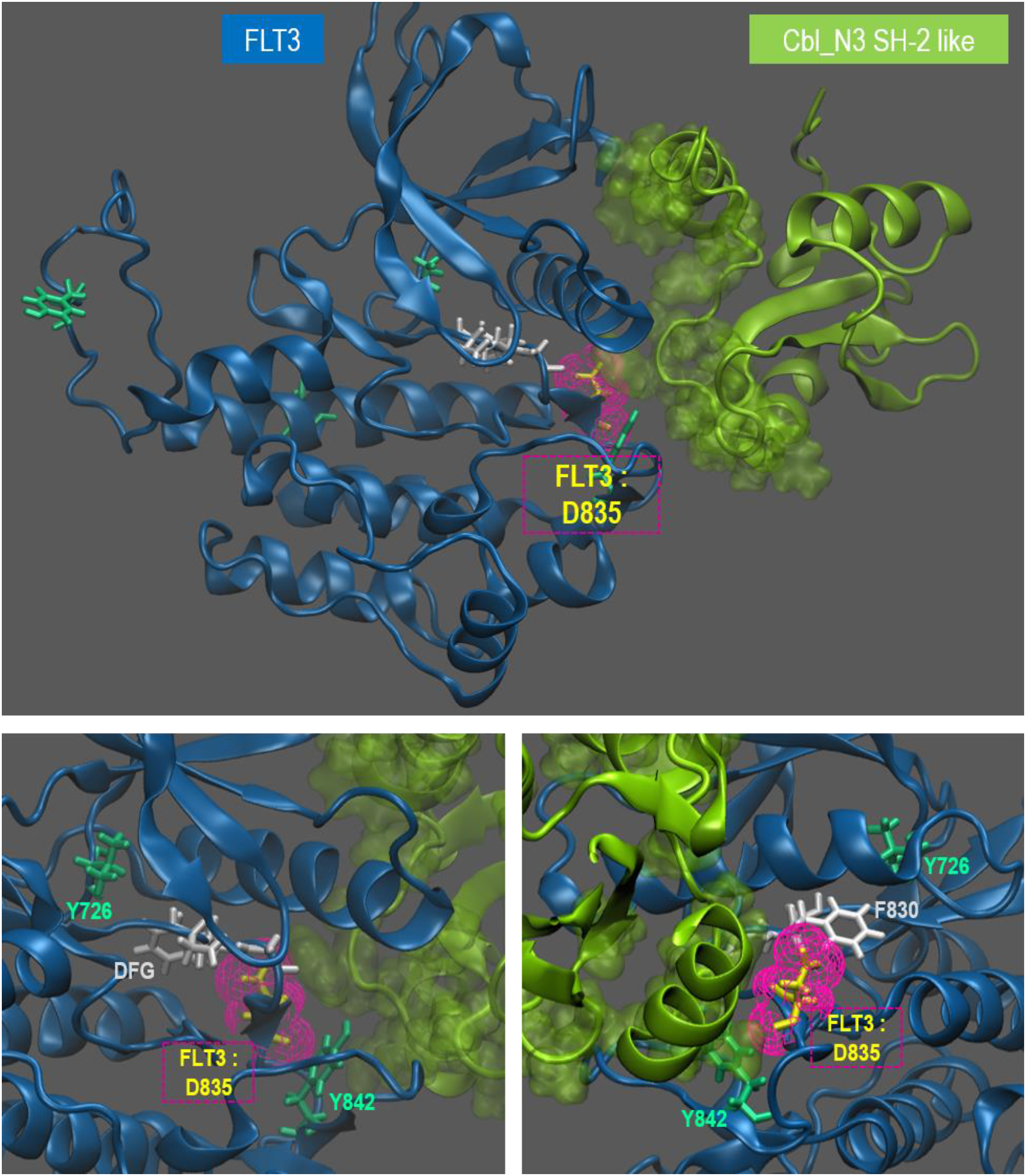
FLT3 and Cbl_N3 protein-protein interaction model. A structural PPI model of active FLT3 kinase and Cbl_N3 SH2-like domains were generated by structural modelling and refinement as described in the pipeline described in the Methods. Briefly, active FLT3 kinase (DFG-IN, αC Helix-IN) and Cbl_N3 domains were modelled for an active KIT kinase (PDB: 1PKG) and whole Cbl_N (PDB: 5HKZ) domains respectively. By applying these modelled structures, two steps of protein interaction prediction were conducted: 1) by PRISM [66] to generate a preliminary interface prediction model and 2) by ROSIE docking [67, 70] to generate multiple refined models from the preliminary model. Docking scores of modelled structures based on contact, van der Waals (VDW), environment and pair-wise interactions between chains were provided and the best scoring one was visualised by VMD software (http://www.ks.uiuc.edu/). Each domain is in a cartoon form, FLT3 (blue) and Cbl_N3 (yellow-green) and the interface of the Cbl_N3 domain in a yellow-green surface form. The FLT3 mutation hotspot, D835 is annotated as sticks inside magenta mesh bubbles. FLT3 tyrosine kinase phosphorylation sites are in green sticks and the DFG(Asp-Phe-Gly) motif is shown as white sticks. (top) whole complex in front; (bottom) zoomed and rotated the y-axis in 25° (left) and −135° (right)

**Figure 8.**
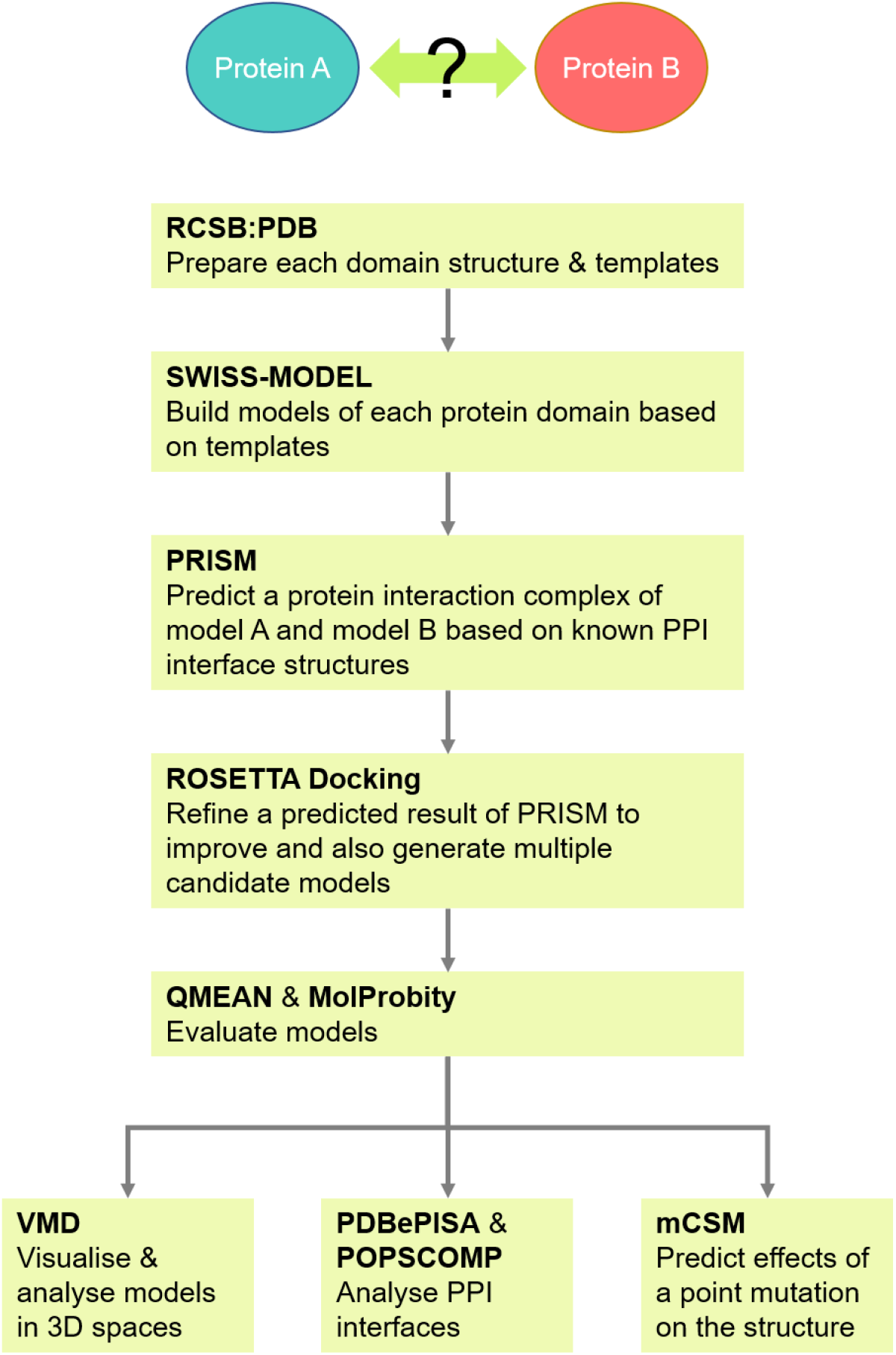
A flowchart of structure modelling to predict a protein-protein interaction complex. This flowchart describes a series of structural modelling applications used for protein-protein interaction prediction.

## Discussion

Proteins and their interactions are essential to form and regulate the complicated machinery of cells in our body. Diseases are outcomes of malfunctions in the molecular mechanisms underlying this machinery. Cancer pathologies are the epitome example showing the complexity of disease mechanisms disrupting cellular functioning with epigenetic and genetic abnormalities changing the expression and functions of the affected proteins. Thus, building a comprehensive protein-protein interaction network of a cell is an important step towards an understanding of molecular disease mechanisms and the development of targeted treatments. Several databases provide amalgamated information on protein interactions data from multiple sources. However, since all of these have incomplete protein-protein interaction network data and are scattered over multiple resources, each with its own format, it is challenging to amalgamate them to form a more complete human proteome map. The UniPPIN presented here, a unified human protein-protein interaction network, integrates multiple resources of human PPIN covering 19,370 proteins with 385,370 interactions. A recent well curated large-scale human protein interaction map integrating high-confidence mass spectrometry experiments covers > 7,700 proteins, and > 56,000 unique interactions [71]. The UniPPIN proteome map is designed to deepen our understanding of human proteomes, as well as the functional relationship between genotypes and phenotypes, especially to extend protein coverage for diseases with largely unknown or understudied molecular mechanisms at the basis of their pathology, such as AML. To explore specific sub-networks efficiently and acquire functional information from the available extensive and scattered data, the short loop network motif profiling method [46] was applied to the amalgamated UniPPIN in the context of Leukemia-specific proteins. We also develop a new concept, that of short loop commonality and propose that protein “modules” that form short loops with different, related proteins may provide selective pressure for a class of hotspot mutations that affect protein-protein interactions.

In our previous study [46], short loops were shown to contain unique information about PPINs. This analysis method demonstrated its utility by retrieving specific and hidden topological and biological properties of the networks. Here we used this approach to analyse PPINs containing mutated proteins in four different leukaemias. The high ratio of short loops in the AML related PPIN shows that mutations reported in AML are more interconnected in the underlying proteome graph than those in other leukaemia PPINs and our analyses indicate that mutated proteins in AML play roles in a broad range of biological processes.

Here, we present a novel approach called ‘short loop commonality’ to analyse indirectly connected proteins having short loop interactions. The method can identify communities of proteins or “modules” related to particular functions. Additionally, we observe the enrichment of interactions between RTK-SH2 domains of these short loop commonality proteins. Indeed, a predicted 3D structural PPI model of active FLT3 kinase and Cbl_N3 SH2-like domain interactions showed the hotspot mutations located at the putative interface of the PPIs involved. This is important as, for example CBL is a ubiquitin ligase that regulates the turnover of its associated RTK [72], and destabilizing the interaction would be predicted to reduce RTK ubiquitination which could prolong the half-life of the RTK at the plasma membrane, affecting signalling pathways. These hotspots can interfere not only with the PPIs involved but also with the phosphorylation of the kinases which could in turn affect degradation of the RTKs involved.

Network biology has improved our understanding of biological systems related to diseases by implementing models based on topological properties of intracellular networks [44, 73, 74]. The aim has been to identify geno-/pheno-typic associations in diseases and ultimately to develop novel translational approaches [73, 74]. Also, recent efforts on human protein interactomes mapped with disease associated proteins have been used to predict how protein abnormalities can affect protein complexes [22, 51]. However, mapping mutation information onto PPINs is still hampered by proteome coverage and the sparsity of information about the sub-networks affected by mutations. The concept of short loop commonality is a promising method for analysing PPINs and finding underlying proteins which affect disease-related mechanisms. It supports a paradigm shift in the drug discovery approach from a one-target-one drug model to a multiple-target strategy [75]. Many mutated proteins are not suitable for drug discovery and we propose that our approach may be useful in identifying functional protein modules affected by short loop commonality mutations that are potential new drug targets. This could provide twofold benefits by tailoring druggable targets to a robust “module” complex, and by predicting drug effects based on similarities of short loop commonality.

With the present study, we confirm that short loop network profiling can be used to analyse genome-wide data of mutations in cancers. This approach may shed light on the underlying functional implications of short-range protein interactions and add useful information about PPINs. Furthermore, proteins sharing short loop interactions may identify essential PPI modules which can affect physiologically important functions and eventually cause cellular diseased states. We propose that this may drive the selection of hotspot mutations. This knowledge can be exploited in the design of experimental targets with measurable phenotypes to understand their mutational effects in cancer or other disease-related mechanisms. These associations will ultimately stimulate the investigation of new protein targets in protein modules of the commonality for drug discovery and drug repurposing.

## Materials and Methods

### Protein-protein interaction datasets

Protein-protein interactions are represented by graph models consisting of nodes of proteins and edges of their interactions. We integrated a data set of 9 different human protein-protein interaction resources including collated databases and recent large-scale studies identifying protein-protein interactions. The studies include a broad binary proteome map by screening pairwise combinations of over 10,000 human open reading frames [76] with yeast-two-hybrid assays [18], collated published evidence (String) [25], affinity purification/mass spectrometry-based networks using different “bait” proteins (green fluorescent protein-tagged (GFP) for [20] and FLAG-HA epitope tags for the BioPlex network [19]) and co-fraction/mass spectrometry-based networks [21, 51]. Protein interaction information from all datasets except for String [25] is derived from direct experimental evidence from the laboratory concerned and only high confidence scored interaction information (above 0.5) is counted from the String database [25]. The UniProt Accession number [77] (collected on March 15^th^ 2017) was used to amalgamate different formats of each dataset and I generated a unified human protein-protein interaction network (UniPPIN) having neither self-loops nor duplicate interactions. The details of the resources are described in Supplementary Table S1.

### Resources of human genetic variations or single nucleotide polymorphisms and mutations in cancer

The Catalogue Of Somatic Mutations In Cancer (COSMIC) [13], the largest cancer mutation database deposited from numerous research institutes worldwide, was used to download cancer and leukaemia related variation information (v80 (Feb 2017)). The database contains information on the somatic mutations present in samples isolated from individual cancer patients. The methods used include whole genome sequencing, exon sequencing, targeted exon/codon sequencing and specific single nucleotide change analyses. Protein mutations reported for several sub-types of four common leukaemias were retrieved: acute myeloid leukaemia (AML), chronic myeloid leukaemia (CML), acute lymphoid leukaemia (ALL) and chronic lymphoid leukaemia (CLL). These four were chosen as they affect different haematopoietic white cell lineages, namely myeloid and T-and B-lymphoid cells. The histology terms and their classification are listed in Supplementary Table S2. Mutation types were selected which result in amino acid changes: substitution nonsense, substitution missense, insertion inframe, insertion frameshift, deletion inframe, deletion frameshift and complex or compound mutation. ‘Whole gene deletion’ and ‘nonstop extension’ mutations are included but both mutations were rarely observed (56 out of 69,334). In addition, only genes with these nonsynonymous mutations in at least two different patients were included. The datasets with ENSEMBL Gene identifiers [78] were mapped into the UniProt Accession number [77] for the analyses in this project.

The leukaemia related mutation datasets were compared with somatic cancer mutation dataset or non-disease human genetic variation information collected from COSMIC and dbSNP of which mutation types are point mutations or single nucleotide variants (SNVs) giving rise to nonsynonymous mutations.

We collected data from different public resources: for disease-associated variant information, nonsynonymous SNVs from COSMIC [13]; for non-disease related information, a subset of dbSNP [30] grouped as common mutations. The details of each dataset and the criteria are: 1) COSMIC exonic variants in variant call format (VCF) (CosmicCodingMuts.vcf) downloaded (v80, February 2017), 2) “common” variants from dbSNP defined in the National Center for Biotechnology Information, U.S. National Library of Medicine (NCBI) database, “germline origin and a minor allele frequency (MAF) ≥ 0.01 in at least one major population, with at least two unrelated individuals having the minor allele”.

These variant datasets in variant call format (VCF) were mapped to the ENSEMBL protein sequences (GRCh37) [78] by using the Variant Effect Predictor (VEP) software tool [79]. The datasets were further filtered for missense variants which map to canonical protein sequences. For each protein, the frequency of localised variants was normalised by the length of amino acid sequences in the protein defined as

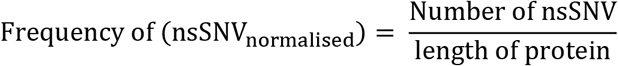

based on the assumption that the mutability of a protein is primarily associated with its size and that differences in amino acid composition between proteins do not have an impact on their overall mutability (nsSNV: nonsynonymous single nucleotide variants).

### Sub-networks of protein-protein interactions related to gene mutations

Proteins of each variant dataset based on the UniProt Accession number (collected on March 15^th^ 2017) were mapped to the UniPPIN. Each dataset of mutated proteins was mapped to the UniPPIN and then mapped proteins and interactions between those proteins were extracted to construct sub-networks related to specific leukaemias or variant datasets. As an example of the labelling used, “AML-related protein-protein interaction network” stands for protein interactions among proteins mutated in at least two AML patients. However, the mutations do not necessarily occur in the same patient.

### Short loop network motif profiling

The short loop network profiling approach [46] was used to analyse PPIN sub-networks containing mutations. The numbers of short loops in each network were calculated and the results were compared with randomised models. The numbers of short loops in the variant specific PPINs and randomly generated PPIN models were evaluated by statistical tests (described below). Functional analyses using Gene Ontology (GO) terms [80] were carried out by measuring functional consensus, i.e. the percentage of GO terms shared by proteins in a short loop as previously described [46]. In addition, g:Profiler [81] and ClueGO [53] were used to measure function enrichment of proteins in different sets. The methods can measure statistical significance of given datasets (p-value ≤ 0.05) compared with their functional term databases (here, Gene Ontology [80]). As there was no significant difference in the topology and ontology of proteins in short loops of different lengths when we applied rigorous graph dynamics and functional enrichment analyses throughout short loop lengths 3 to 6 for the previously studied larger network [46], only short loop interactions with length 3 were used in this study.

### Structural modelling, refinement and model evaluation

We focused our structural studies on the protein with the most frequent pathogenic mutations Fms Related Tyrosine Kinase 3 (FLT3), other tyrosine kinase proteins sharing ‘short loop commonality with FLT3 (described in Figure 1) and the other short loop interactions. We hypothesised that “active” tyrosine kinases interact with SH2 domains *via* the middle of the kinase domain between N-and C-lobes. To test this, we built a three-dimensional protein-protein interaction model of active FLT3 and Cbl_N3 SH2-like domains by using a pipeline described in Figure 8. We retrieved different 3D structures of FLT3, KIT and CBL functioning ubiquitination of these kinases as reference templates and used a series of publicly available web applications involving SWISS-MODEL to build a single protein structure complex based on the templates (https://swissmodel.expasy.org/) [82], PRISM for predicting protein-protein interaction interfaces by structural matching (http://cosbi.ku.edu.tr/prism/) [66, 83] and the RosettaCommons(https://www.rosettacommons.org/) docking webserver called ROSIE (http://rosie.rosettacommons.org/docking2) [70] for refining the predicted model from PRISM. To evaluate the quality of the models, MolProbity (http://molprobity.biochem.duke.edu/) [84] and QMEAN (https://swissmodel.expasy.org/qmean/) [85] scores were used. Protein-protein interaction interfaces were analysed by PDBePISA (http://www.ebi.ac.uk/pdbe/pisa/)[86] and POPSCOMP (https://mathbio.crick.ac.uk/wiki/POPSCOMP) [87]. For visualization of the structures, structural alignments and analysis, the VMD software (http://www.ks.uiuc.edu/) was used.

## Acknowledgements

This research was supported by Bloodwise (to NSBT, FF and SSC) and the British Heart Foundation (RE/13/2/30182 to FF and AL).

